# A feedback between dispersal and hybridization may facilitate the repeated emergence of flight in field crickets

**DOI:** 10.64898/2026.09.21.753268

**Authors:** Leonardo T. Gonçalves, David A. Gray, Lisa A. Treidel, Colin D. Meiklejohn, Kristi L. Montooth

## Abstract

Dispersal and hybridization can reinforce one another in a feedback loop when species that disperse farther are more likely to encounter and mate with close relatives, resulting in gene flow that can reintroduce dispersal-enhancing alleles in their descendants. This dynamic could help explain why complex dispersal-related traits, such as flight in insects, appear to evolve repeatedly within a clade, but this idea has rarely been tested with genomic data. North American *Gryllus* field crickets, in which flight capability appears to have been regained from flightless ancestors at least nine independent times, offer a powerful system to do so. Using a genome-wide dataset, we test three predictions about the origins of phylogenetic discordance in *Gryllus* and the role of dispersal-related traits. First, we find that both incomplete lineage sorting (ILS) and hybridization contribute to extensive gene tree conflict observed across the *Gryllus* phylogeny, but several conflicting patterns cannot be explained by ILS alone. Multiple lines of evidence reveal introgression at both deep and recent nodes, producing a mosaic evolutionary history with reticulation across many lineages. Second, hybridization is disproportionately concentrated among flight-capable lineages, supporting the idea that dispersal ability increases opportunities for interspecific gene flow. Third, introgression from flight-capable lineages could help explain repeated trait reversal across the phylogeny, suggesting that allele flow may have contributed to the re-emergence of flight in descendants of flightless ancestors. These results are consistent with dispersal-hybridization feedback shaping both the reticulated phylogenetic history of *Gryllus* and the repeated emergence of flight. More broadly, this species group reveals how models of character evolution built on bifurcating trees may overestimate the number of independent trait origins when reticulate processes are ignored.

**Teaser text:** Evolutionary histories are often networks rather than bifurcating trees, with important consequences for how we interpret repeated trait evolution. In North American field crickets, we show that both incomplete lineage sorting and hybridization contribute to extensive discordance among gene trees, with introgression detectable across deep and recent timescales. Hybridization disproportionately involves flight-capable lineages and several introgression events coincide with lineages that regained flight from flightless ancestors. This supports a feedback loop, in which dispersal promotes hybridization, and hybridization in turn reintroduces genetic variation that can restore flight capability. More broadly, our results suggest that repeated origins of complex traits inferred from bifurcating phylogenies may instead partly reflect reticulate evolutionary histories.

## Introduction

Recent advances in phylogenomics are reshaping our understanding of evolutionary relationships, challenging the classical bifurcating “Tree of Life” model. In many plant and animal groups, gene trees inferred from independent loci show extensive discordance with one another and with species-level phylogenies (e.g., Pease *et al*., 2016; Pezzi *et al*., 2024; Mars *et al*., 2025; Thomas *et al*., 2025). Such conflicts may arise from incomplete lineage sorting (ILS)—the persistence of ancestral polymorphisms across speciation events—, but can also reflect hybridization and introgression, in which genetic material moves across species boundaries through gene flow (Maddison, 1997). Whereas ILS is expected in rapid radiations with large ancestral population sizes, the probability of introgression is influenced by the propensity of disparate populations or species to hybridize and the selective effects of foreign alleles in recipient populations. Introgression is therefore more likely than ILS to alter evolutionary trajectories by transferring adaptive alleles, reintroducing ancestral variation, or homogenizing species boundaries (Cui *et al*., 2013; Harrison & Larson, 2014; Uckele *et al*., 2024). Together, these processes contribute to a network-like history of life, where reticulation rather than strict divergence increasingly appears to be the rule (Mallet *et al*., 2016). Disentangling ILS from introgression is possible due to the directional signal that introgression leaves in the genome. Yet, it remains a central challenge when both processes act simultaneously, particularly when there is uncertainty in population sizes and species divergence times.

Dispersal ability is a key, and often underappreciated, driver of how frequently hybridization occurs between species. Individuals capable of long distance movement are more likely to encounter members of other species, cross geographic barriers that otherwise would maintain reproductive isolation, and establish contact zones where gene flow becomes possible (Lowe *et al*., 2015; Waters *et al*., 2020). This relationship between dispersal and hybridization may generate a positive feedback loop, where dispersal-capable individuals hybridize more frequently, and if hybrid or admixed offspring inherit or retain traits that enhance dispersal, they in turn disperse further and encounter additional heterospecifics, accelerating the spread of introgression through space—a process Shine et al. (2011) termed spatial sorting. This feedback is particularly tractable in wing-dimorphic insects, because variable dispersal capability is encoded as a visible, binary, and often heritable polymorphism. Fully winged, flight-capable morphs are overrepresented at range expansion fronts (Simmons & Thomas, 2004), where encounters with other species are most likely, and where hybridization rates may thus be elevated relative to populations at the center of the range. If introgression between flight-capable and flightless lineages reintroduces alleles associated with flight, selection at range expansion fronts could sustain flight polymorphisms across lineages that would otherwise lose them, and even facilitate the recovery of flight capability in lineages where it had been reduced or lost. This positive feedback between dispersal and hybridization has broad implications for understanding both the dynamics of gene flow and the evolution of traits that influence dispersal itself, including whether alleles enabling flight can spread across lineage boundaries through introgression.

North American *Gryllus* field crickets offer an ideal system to test these ideas. The clade encompasses 35 valid species and seven lineages that may represent undescribed species (Weissman & Gray, 2019; Gray *et al*., 2020) and has served as a model system at the intersection of evolution, behavior, physiology, and ecology. It represents a recent radiation, supported by the conserved morphology across species (Gorochov, 2019) and by estimates that most diversification occurred within the last 1 mya (Gray *et al*., 2020). Several instances of hybridization are documented between species pairs, both in controlled laboratory crosses and in natural populations (Gray, 2005; Izzo & Gray, 2011; Larson *et al*., 2013; Veen *et al*., 2013; Gray *et al*., 2016; Panagiotopoulou *et al*., 2016), with the hybrid zone between *G. firmus* and *G. pennsylvanicus* in the eastern United States representing one of the best-studied examples of reproductive isolation and gene flow in insects (Ross & Harrison, 2002; Larson *et al*., 2013; Grapin *et al*., 2026). Beyond hybridization, most *Gryllus* species have large effective population sizes that increase the potential for ILS (Broughton & Harrison, 2003; Gray *et al*., 2020), which further complicates phylogenetic inference. Gray et al. (2020) produced a comprehensive molecular phylogeny of North American *Gryllus* using Anchored Hybrid Enrichment (AHE; Lemmon et al. 2012) and noted that both ILS and hybridization were likely contributing to observed gene tree conflict. However, the relative contributions of these processes remain untested along with their implications for trait evolution.

A robust phylogenetic framework that accounts for reticulate evolution may be critical for understanding the repeated emergence of flight-capability in *Gryllus*. North American *Gryllus* includes monomorphic long-winged (LW; flight-capable) and short-winged (SW; flightless) species, as well as wing dimorphic (WD) species in which both morphs occur within populations (Figure 1B). Wing dimorphism in crickets represents a classic life history trade-off: SW morphs are flightless but invest more in early reproduction, while LW morphs disperse at a cost to fecundity (Harrison, 1980; Roff, 1986; Zera & Denno, 1997). Ancestral character reconstruction suggests that WD lineages have evolved from flightless SW ancestors nine independent times, suggesting that wing dimorphism is not simply a transitional state between a fully winged ancestor and permanent loss of flight (Treidel *et al*., 2026). However, these repeated transitions remain controversial, as they may reflect artifacts of model-based trait reconstruction or unaccounted phylogenetic uncertainty. Given the shallow divergence times among *Gryllus* species and the polygenic threshold-trait basis of wing-morph determination (Zera & Rankin, 1989; Roff & Fairbairn, 1991), it is plausible that alleles promoting flight capability have been maintained as standing variation in flightless lineages, or have entered these lineages through hybridization with flight-capable relatives (Harrison & Larson, 2014; Waters *et al*., 2020). If flight itself promotes the dispersal and secondary contact that make hybridization more likely, the repeated emergence of flight and the reticulate history of the clade may not be independent.

**Figure 1.**
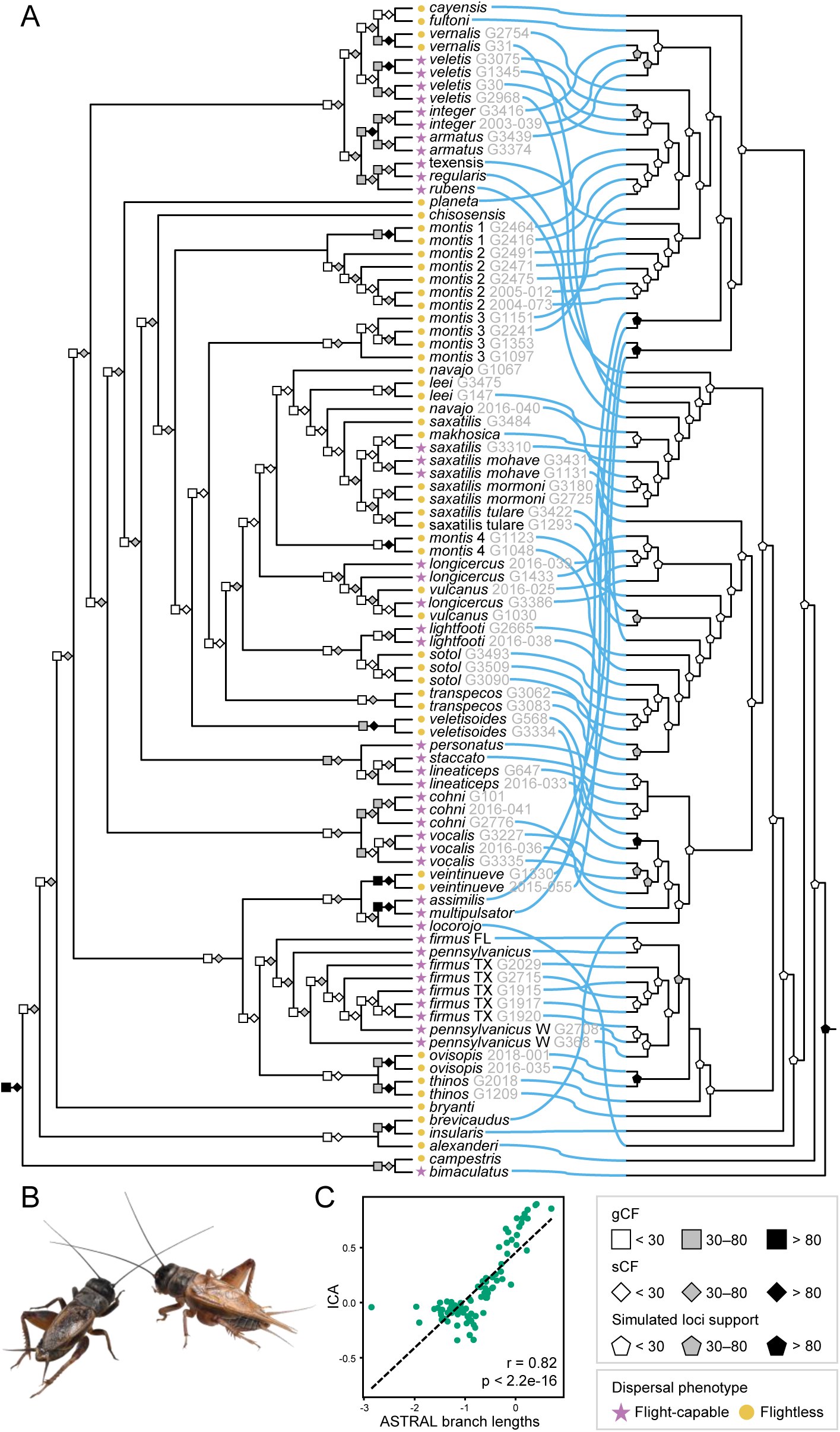
Phylogenetic discordance across genomic compartments and sources of gene tree conflict in North American *Gryllus*. (A) Tanglegram comparing the ASTRAL coalescent species tree (left) and the mitochondrial maximum likelihood tree (right). Connecting lines link the same taxon across trees. Nodes on the ASTRAL tree are annotated with Gene Concordance Factors (gCF) and Site Concordance Factors (sCF). Nodes on the mitochondrial tree are annotated with the percentage of nuclear gene trees simulated under the ASTRAL topology that recovered that mitochondrial clade (see Methods). Tip symbols indicate dispersal phenotype. Outgroups were pruned for visualization. (B) Representative photographs of the short-winged (flightless, left) and long-winged (flight-capable, right) morphs of *Gryllus lineaticeps*, a wing dimorphic species included in this study. Wing dimorphic lineages were classified as flight-capable throughout all analyses. Photo credit: Essig Museum of Entomology. (C) Relationship between branch lengths in the ASTRAL species tree (in coalescent units, log_10_ transformed) and Internode Certainty All (ICA) values.

Here, we reexamine the evolutionary relationships among North American *Gryllus* species using sequence data generated by Gray et al. (2020), supplemented with mitochondrial genomes we assembled from off-target AHE reads. We leverage these data to test the relative contributions of ILS and hybridization to phylogenetic discordance, and to evaluate whether reticulate evolution may have contributed to the repeated transitions in wing morphology. Specifically, we tested the following hypotheses: (i) hybridization across both recent and deep evolutionary history in *Gryllus* contributes to phylogenetic discordance reported in previous works; (ii) flight-capable lineages (WD and LW) are disproportionally involved in introgression events, consistent with the prediction that flight promotes dispersal, secondary contact, and interspecific gene flow; and (iii) repeated independent origins of flight from flightless ancestors may partly reflect introgression from flight-capable lineages rather than independent origins, potentially confounding traditional models of character state evolution that assume a strictly bifurcating history.

## Material and Methods

### Dataset

We leveraged the AHE dataset published by Gray et al. (2020), which was used to infer the first comprehensive molecular phylogeny of North American *Gryllus* crickets. The dataset included representatives of all 35 described species, as well as seven additional lineages that may represent undescribed taxa. The ingroup comprised 89 individuals, with most species sampled multiple times (Table S1). Two Old World *Gryllus* species (*G. campestris* and *G. bimaculatus*) and three outgroup species from other genera within Gryllidae (*Acheta domesticus*, *Nirgogryllus sibiricus*, *Teleogryllus emma*) were also included.

The dataset spans 560 nuclear loci, totaling 492,531 aligned nucleotide sites. On average, each individual is represented in 530 of the 560 loci. To maintain consistency with the original study, we used the alignments provided by Gray et al. (2020) without modification. For downstream analyses, we split the concatenated alignment into individual loci using Biostrings (Pagès *et al*., 2023), guided by the partition file provided by the authors. Although Gray et al. (2020) reported a total of 563 loci in their dataset, only 560 loci were recovered using the partition file, likely reflecting minor filtering steps that were not explicitly documented.

To gain additional insights into phylogenetic conflict, we also assembled mitochondrial loci from off-target AHE data. Reads were trimmed and filtered with fastp v0.23 (Chen *et al*., 2018) and then mapped against the *G. veletis* Refseq mitochondrial genome (NC_057053.1; Torson et al. 2022) using BWA v0.7.17 (Li, 2013). Consensus sequences were generated with samtools v1.20 (Li *et al*., 2009), using default settings and a minimum coverage of 4x, following Lyra et al. (2017). Protein-coding and ribosomal genes were manually extracted by aligning consensus mitogenomes against the reference genome with MAFFT v7 (Katoh *et al*., 2019). The alignment was manually inspected using AliView v1.28 (Larsson, 2014), and orthology was assigned based on reference genome coordinates. Extracted loci were concatenated into a supermatrix, and samples with ≥50% missing data were excluded from downstream analyses.

Trait data on wing morphs were obtained from Treidel et al. (2026), which was built upon extensive field records of adult morphology (Weissman & Gray, 2019). Species were classified as flight-capable (LW, WD) or flightless (SW). These classifications were used in downstream comparative analyses testing the relationship of dispersal phenotype with admixture signals.

### Phylogenetic inference and analyses of tree discordance

We employed two methods to infer phylogenetic relationships. First, a maximum likelihood tree was inferred in IQ-TREE v2.3.4 (Minh *et al*., 2020) from the concatenated alignment (supermatrix approach), assuming a GTR+Γ model of sequence evolution for each gene partition and testing for node supports using ultrafast 1,000 bootstrap replicates. Second, we inferred gene trees individually in IQ-TREE under the same model and bootstrap scheme. Nodes with ≤10% support were collapsed using Newick Utilities v1.6 (Junier & Zdobnov, 2010). These gene trees were then used as input to construct a quartet-based species tree consistent with the multispecies coalescent model (supertree approach) using ASTRAL-III v5.7.8 (Zhang *et al*., 2018).

To assess phylogenetic conflict, we calculated gene concordance factors (gCF) and site concordance factors (sCF) in IQ-TREE (Mo *et al*., 2023), using the ASTRAL topology as reference. We selected the coalescent tree as reference because it explicitly accounts for ILS and integrates information across loci, providing a more appropriate baseline for conflict quantification (Degnan & Rosenberg, 2009; Mirarab *et al*., 2014). We also applied Quartet Sampling (QS) (Pease *et al*., 2018) to the same topology, which evaluates localized support and discordance among quartets. Following Pease *et al*. (2018), we interpret quartet concordance (QC) values above 0.2 as strong support for the reference topology, between 0 and 0.2 as indicating conflict among gene trees, and below –0.2 indicating strong support for an alternative relationship. Quartet differential (QD) will be 1 when the two non-focal topologies are recovered at equal frequency, as expected under ILS, especially when QC indicates conflict with the first quartet. When conflict is asymmetrically biased towards one alternative topology, as expected under introgression, QD will approach zero. In addition, we calculated Internode Certainty All (ICA) scores with RAxML v8.2.12 (Stamatakis, 2014), which summarize the frequency of alternative bipartitions across gene trees. ICA values close to 1 indicate strong concordance for the bipartition defined by a given internode, while ICA values close to 0 indicate strong conflict. Negative ICA values indicate that the defined bipartition conflict with other high frequent bipartitions (Salichos *et al*., 2014). We examined the correlation between ASTRAL tree branch lengths and ICA values using Pearson’s correlation in R v4.3.2 (R Core Team, 2023), testing whether shorter coalescent times are associated with stronger conflict, which is likely explained by ILS (Zhou *et al*., 2022).

Mitonuclear discordance was investigated by comparing the mitochondrial supermatrix tree with the nuclear ASTRAL tree. The mitochondrial tree was inferred in IQ-TREE using the GTR+Γ model with 1,000 ultrafast bootstraps. To test whether discordance could be explained by ILS alone, we simulated 5,000 nuclear gene trees under the ASTRAL topology using the function sim.coal.mpest in the R package Phybase v2.0 (Liu & Yu, 2010). Simulated trees were used to calculate gCF onto the mitochondrial topology with IQ-TREE. If ILS alone explains discordance, mitochondrial clades should be present among simulated nuclear gene trees. Conversely, absence or low frequency of mitochondrial clades on simulated nuclear gene trees, would be more consistent with introgression (Rose *et al*., 2025).

### Assessing introgression and an association with flight-capability

We applied multiple approaches to test for introgression. First, we conducted rooted triplet (RT) tests (Larson *et al*., 2021) on multiple three taxa subsets from the phylogeny. This test compares the proportions of loci that support each of the three possible topologies for a given set of three taxa and an outgroup. Under the multispecies coalescent model, the most common (major) topology is expected to reflect the true species relationship, and the two less common (minor) topologies are expected to result from ILS. Assuming a null hypothesis of ILS only and no admixture, the two minor relationships should be equally frequent within the distribution of gene trees, and significant deviations from equality may be considered evidence for gene flow (Pamilo & Nei, 1988; Degnan & Rosenberg, 2009; Hibbins & Hahn, 2022). To generate the set of triplets to test, we sampled each pair of sister species from the tree and paired them with each of the remaining samples, resulting in 705 input tests. In cases where one of those lineages was poly– or paraphyletic in the species tree, additional representatives were included to capture the observed splits. For each RT, gene alignments were pruned to include only the three ingroup taxa of interest and the outgroup *A. domesticus*. Then, gene trees were estimated with IQ-TREE. We compared the observed distributions of minor topologies to a binomial distribution with the probability of either minor relationship being equal. P-values were adjusted for multiple comparisons with the False Discovery Rate method (Benjamini & Hochberg, 1995) and a significance threshold of 0.05. RT tests were conducted with published Python scripts (Larson *et al*., 2021) that use IQ-TREE and phyx (Brown *et al*., 2017) as dependencies.

In addition to the RT tests, we used HyDe v1.0.2 (Blischak *et al*., 2018) to evaluate evidence of hybridization based on phylogenetic invariants. HyDe estimates an admixture parameter (γ), where values near 0.5 indicate equal genomic contributions from parental species, and values approaching 0 or 1 reflect asymmetrical contributions. Individuals were assigned to species, resulting in 55,272 trios. To test whether hybridization is associated with flight capability, we coded taxa by their dispersal phenotype: flightless (SW) versus flight-capable (LW, WD). We then used a permutation test in R to test whether flight-capable species were overrepresented in significant HyDe triplets. Flight capability labels were randomly reassigned among taxa 10,000 times, and for each iteration we computed the proportion of significant triplets involving at least one flight-capable species. The observed proportion was compared to this null distribution to obtain an empirical p-value.

Finally, we applied a model-based phylogenetic network approach to distinguish between recent and ancient hybridization. We used SNaQ (Solís-Lemus & Ané, 2016) to infer networks that accommodate reticulation, testing hypotheses of zero up to four reticulation events (h). Concordance factors were estimated from gene trees with readTrees2CF implemented in PhyloNetworks v1.2.0 (Solís-Lemus *et al*., 2017). The ASTRAL tree served as the starting topology for the h = 0 search, and the best network from each iteration was used as the starting topology for the subsequent search. Because of the computational burden of these analyses, we reduced the dataset to representatives of the major *Gryllus* lineages identified by Weissman and Gray (2019), selecting individuals with the greatest locus recovery.

## Results

### Rampant gene tree discordance in the field cricket radiation

We reanalyzed the Gray et al. (2020) AHE dataset using both supertree (ASTRAL) and supermatrix (IQ-TREE) approaches to reconstruct nuclear phylogenies. Both methods recovered topologies concordant with the original study, with high support for most branches (LPP = 0.95– 1 in ASTRAL and BS = 100 in IQ-TREE; Figure S1). Concordance factor statistics (gCF, sCF) revealed that, despite high support values, only a small fraction of individual gene trees and sites support several backbone nodes (Figures 1A, S2 and S3). Moreover, no single gene tree mirrored the species tree topology. Such discordance is indicative of a history of introgression and/or ILS. Quartet-based evaluation of the ASTRAL tree found that, on average, a third of quartets were informative (QI = 0.34; Figure S4), indicating that the data carry moderate signal for resolving relationships at many nodes. Quartet concordance was very low on the backbone (average QC = 0.04) and within intraspecific clades (average QC = 0.07), but substantially higher among shallow, sister species relationships (average QC = 0.44). Differential quartets were frequently asymmetric (QD values < 1; Figure S4), indicating that conflict often favored one of the two alternative topologies, a pattern expected under introgression rather than ILS (Pease *et al*., 2018).

ICA values, which were low for backbone and intraspecific nodes, were consistent with these results and reflected conflict among the relationships (Figure S5). Finally, ASTRAL branch lengths were positively correlated with ICA (Pearson’s r = 0.82, p < 2.2e-16; Figure 1C), consistent with ILS contributing disproportionately to conflicts among short coalescent intervals (Zhou *et al*., 2022; Jin *et al*., 2025).

### Mismatches between nuclear and mitochondrial topologies

We recovered off-target mitochondrial sequences from the AHE raw data to evaluate mitonuclear concordance (Table S1). The mitochondrial phylogeny conflicted with the ASTRAL tree in several nodes, revealing extensive mitonuclear discordance (Figures 1A and S6). For example, *G. montis* lineage 2 grouped with *G. montis* lineage 1 in the nuclear tree but with *G. texensis* in the mitochondrial tree. Several species, including *G. thinos* and *G. lightfooti*, were para– or polyphyletic in the mitochondrial phylogeny despite nuclear monophyly. Simulations of nuclear gene trees under the ASTRAL species tree revealed that deep mitochondrial nodes were rarely recovered (Figure 1A), suggesting that ILS alone cannot explain the mitochondrial backbone and that historical introgression may have contributed to mitonuclear discordance. At shallower nodes, simulated nuclear trees matched mitochondrial clades more frequently, consistent with ILS, but some shallow paraphyletic clusters occurred less frequently than expected, implicating localized mitochondrial introgression. These instances often involved species with documented natural hybridization, including *G. firmus* × *G. pennsylvanicus* (Ross & Harrison, 2002; Larson *et al*., 2013) and *G. lineaticeps* × *G. staccato* (Gray et al. 2016).

### Introgression events more often involve flight-capable lineages

To test whether there is a relationship between dispersal ability and interspecific hybridization, and to evaluate whether introgression may have contributed to repeated evolutionary transitions in flight capability, we quantified patterns of gene flow across the North American *Gryllus* radiation using complementary methods.

Using RT tests, we identified 50 species pairs showing significant evidence of introgression (FDR < 0.05; Figure 2). When partitioned by dispersal phenotype, 84% of these inferred introgression events involved at least one flight-capable species, despite the predominance of flightless lineages in the dataset (Figure 2). This asymmetry supports the prediction that enhanced dispersal increased opportunities for interspecific contact and gene flow. Notably, most inferred events (58%) occurred between a flight-capable and a flightless lineage, indicating that introgression frequently bridges contrasting dispersal phenotypes. When mapped onto the species phylogeny, six of these signals coincide with lineages inferred to have independently regained flight (*G. lightfooti*, *G. longicercus*, *G. staccato*, *G. saxatilis, G. saxatilis mohave*, and *G. vocalis*; Figure 2; Treidel *et al*., 2026), suggesting a potential link between introgression and repeated gains of flight.

**Figure 2.**
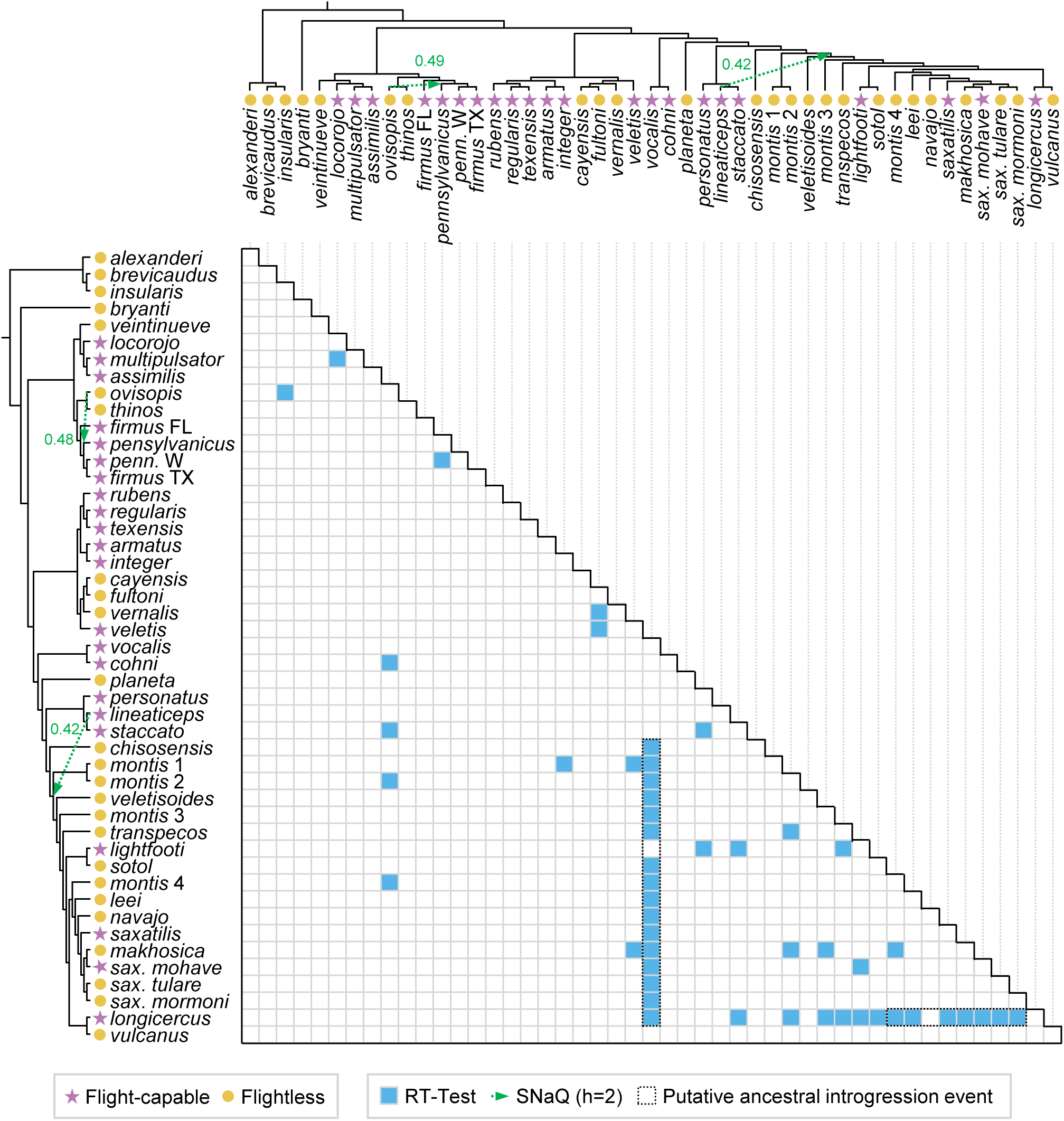
Phylogenetic context of rooted triplet (RT) test results and network inference across *Gryllus* species. Rows and columns correspond to lineages ordered according to the ASTRAL coalescent species tree, shown along the left and top margins. Blue cells indicate species pairs for which the RT test detected a significant excess of gene trees supporting a minor topology inconsistent with the species tree relationship; white cells indicate non-significant comparisons. Tip symbols indicate dispersal phenotype (purple stars, flight-capable; yellow circles, flightless). Green arrows overlaid on the phylogenies indicate the two inferred reticulation edges from the best-fit SNaQ network (h = 2). Blocks outlined in black highlight cases of putative ancestral introgression, in which a single lineage shows significant RT test signals with multiple members of a clade, consistent with a single ancient hybridization event rather than multiple independent recent events.

To explore whether ancestral introgression could have contributed to these transitions, we reconstructed phylogenetic networks using SNaQ on a reduced dataset. The best-fitting network inferred two major hybridization events (h = 2; Figure S7). One involved a substantial genomic contribution (42%) from the ancestor or an unsampled sister lineage of the flight-capable *G. lineaticeps* into the ancestor of a predominantly flightless clade that later experienced three independent reemergences of flight (Figure 2; Treidel et al. 2026). A second event inferred a comparable contribution (49%) from the flightless *G. ovisopis* into the flight-capable *G. pennsylvanicus*, indicating that directionality of introgression is not restricted to flight-capable taxa. Together, these network results are consistent with a history of gene flow both preceding and following transitions in dispersal capability.

We then tested whether hybridization signals were associated with dispersal phenotype by comparing the observed proportion of significant HyDe triplets involving at least one flight-capable parental lineage to a null expectation generated by permutation. To first examine genome-wide introgression, we performed HyDe analyses across all possible lineage triplets, yielding 55,272 tests of putative hybrid ancestry. Of these, 4,912 triplets (8.9%) showed significant evidence of hybridization. The estimated admixture parameter (γ) exhibited a unimodal distribution centered on intermediate values (0.3–0.7; Figure S8A), consistent with widespread and relatively symmetric gene flow rather than highly directional introgression. Among significant triplets, 78.1% involved at least one flight-capable parental lineage, exceeding the null expectation based on random assignment of flight capability (mean = 72.4%, 95% CI = 71.2–73.6%; permutation test, p < 10e-05; Figure S8B). This enrichment indicates that flight-capable species are disproportionately represented in inferred introgression events, consistent with the hypothesis that increased dispersal ability enhances opportunities for interspecific encounters and gene flow. Collectively, these results suggest that introgression may be an important process accompanying repeated evolutionary transitions in dispersal capability within North American *Gryllus*.

## Discussion

Rapid radiations with large ancestral effective population sizes are expected to produce pervasive gene tree discordance, and North American *Gryllus* field crickets are no exception. Our analyses support a scenario in which rapid diversification generated a baseline of discordance driven by ILS that was further complicated, at both recent and deep timescales, by interspecific gene flow. Nodes accompanied by shorter coalescent intervals accumulated disproportionate conflict across gene trees, confirming that ILS is a major contributor to observed discordance. Nevertheless, several patterns in our data cannot be explained by ILS alone and suggest a history of hybridization operating alongside it. Low QC and asymmetric QD scores at multiple nodes indicate that gene tree conflict is biased toward one alternative topology rather than being distributed equally between the two, as predicted under a purely ILS model (Pease *et al*., 2018). Similarly, our coalescent simulations show that deep mitochondrial clades are rarely recovered among nuclear gene trees simulated under the species tree topology, suggesting that mitochondrial genealogy has been shaped by processes, most parsimoniously introgression, that lie outside what ILS alone predicts. These results are consistent with the high levels of gene tree conflict previously reported for the clade (Gray *et al*., 2020), but now with quantified contributions from each process.

The independent lines of evidence for introgression suggest that interspecific gene flow was pervasive during the diversification of the clade. We identified 50 species pairs where gene tree asymmetry exceeded ILS expectations, while the phylogenetic network placed two deep reticulation events with large genomic contributions (42% and 49%) that suggest substantial admixture. Moreover, the unimodal distribution of γ values from HyDe, centered on intermediate values, is consistent with widespread hybridization involving roughly balanced parental contributions across several lineages, a pattern reported in other fast diversifying clades with porous species boundaries (Blischak *et al*., 2018; Edelman *et al*., 2019). This is biologically plausible given that reproductive isolation in *Gryllus* appears incomplete across multiple recently derived species pairs. We note, however, that the AHE probe design targets conserved exonic regions, which may reduce power to detect introgression and affect estimates of gene flow since these loci are a biased subsample of the genome. Future work based on population whole-genome resequencing would allow a more complete characterization of admixture across the clade, including identification of genomic regions that are either permeable or resistant to gene flow.

Our prediction that flight capability promotes interspecific gene flow by enabling dispersal and secondary contact received support across multiple analyses. The overrepresentation of flight-capable lineages among significant introgression signals is congruent with the idea that dispersal increases encounter rates with heterospecifics and therefore the probability of interspecific mating. Interestingly, all *Gryllus* hybrid zones documented in natural populations involve at least one flight-capable species. The link between dispersal ability and hybridization has been proposed broadly for insects (Waters *et al*., 2020), but to our knowledge this is the first genome-wide test of the prediction across a clade. Alternatively, these results may reflect the fact that flightless, SW monomorphic species tend to occupy smaller and more geographically restricted ranges than flight-capable species (Weissman & Gray, 2019), and may therefore have fewer opportunities for secondary contact regardless of dispersal potential.

Our data point to a potential contribution of reticulate evolution to the repeated emergence of flight capability in the North American *Gryllus* phylogeny. Ancestral character reconstruction placed the repeated origins of flight from flightless ancestors at nine independent nodes (Treidel *et al*., 2026), a pattern that challenges the idea that complex traits are not regained once they are lost (Elmer & Clobert, 2025). Such apparent reversals in stick insects have provoked debate, and repeated wing gains from wingless ancestors were later reattributed, at least in part, to retention of developmental potential and shared standing genetic variation rather than true *de novo* evolution (Whiting *et al*., 2003; reviewed in Forni *et al*., 2026). Similarly, alleles underlying flight capability may never have been fully purged in lineages inferred as flightless, either because selection against them was weak in smaller isolated populations, or because their pleiotropic roles in other traits maintained them. For instance, juvenile hormone is known to affect both wing morphology and reproduction in cricket species (Zera & Rankin, 1989; Roff & Fairbairn, 1991). Second, ancestral state reconstruction is itself sensitive to the underlying tree topology (Mooers, 2004; Duchêne & Lanfear, 2015), and the reticulate history we document here means that any bifurcating tree used as reconstruction input may misplace transitions, potentially inflating the apparent count of independent origins.

With these alternatives in mind, we found that introgression provides a plausible complementary mechanism for at least part of the observed transitions. Six lineages inferred to have independently regained flight (*G. lightfooti*, *G. longicercus*, *G. staccato*, *G. saxatilis, G. saxatilis mohave,* and *G. vocalis*) show introgression with flight-capable relatives, and the best-fitting network places a major admixture event (42% genomic contribution) from the branch of the flight-capable *G. lineaticeps* into the ancestor of a clade that subsequently experienced multiple regains of flight. If transferred genomic material included alleles contributing to the developmental switch between wing morphs, then introgression may have facilitated re-emergence of flight in descendant lineages. This mechanism parallels hypotheses for how introgression facilitated repeated evolution of mimicry and visual preference in *Heliconius* butterflies (Heliconius Genome Consortium, 2012; Wallbank *et al*., 2016; Rossi *et al*., 2024) and convergent adaptation in African cichlids (Koblmüller *et al*., 2007; Gante *et al*., 2016; Hulsey *et al*., 2018), where pre-assembled genetic solutions moved across species boundaries rather than being rebuilt from scratch. We emphasize, however, that the overlap between introgression signals and the regain of flight, while suggestive, is not sufficient evidence for a causal link. A definitive test of the introgression hypothesis will ultimately require identifying the specific loci underlying wing morph determination across multiple *Gryllus* lineages and asking whether lineages inferred to have regained flight through introgression share alleles at those loci.

Several additional sources of uncertainty are worth noting. Wing-morph frequency data in *Gryllus* are often heavily skewed, with minority morphs comprising a small fraction of sampled individuals and sometimes going entirely undetected in poorly sampled species (Treidel *et al*., 2026). This means some species classified as flightless (SW) in our dataset may in fact be dimorphic, which would affect our classification of species as flight-capable or flightless when interpreting the introgression analyses. Our dataset is also restricted to North American *Gryllus*, and whether the patterns of reticulate evolution and the association between dispersal and hybridization documented here extend to other *Gryllus* clades or to other wing dimorphic insect groups remains an open question.

Taken together, our results suggest that the evolutionary history of North American *Gryllus* is shaped by an interplay of ILS arising from rapid diversification, reticulate gene flow facilitated in part by flight capability, and positive feedback between dispersal and hybridization that may have left signatures on both the phylogeny and the repeated emergence of flight. These findings contribute to growing evidence that hybridization is a recurrent rather than exceptional process in animal radiations (Seehausen, 2004), and highlight that ancestral state reconstruction on bifurcating trees will miscount trait origins when the true history of a clade is reticulate, a methodological concern that extends beyond *Gryllus* (Hibbins *et al*., 2020). As genomic datasets increasingly reveal network-like histories across the tree of life, integrating reticulate evolution into models of phenotypic diversification will be essential for interpreting patterns of convergence and parallelism.

## Data and code availability statement

Raw reads are available at NCBI under BioProject PRJNA1509682. Data and code supporting this manuscript are available on Dryad: <link>.

## Author contributions

Conceptualization: LTG, CDM, KLM; Methodology: LTG, DAG; Investigation: LTG; Formal analysis: LTG; Visualization: LTG; Funding acquisition: CDM, KLM; Writing—original draft: LTG; Writing—review & editing: LTG, DAG, LAT, CDM, KLM.

## Acknowledgements

This work was completed utilizing the Holland Computing Center of the University of Nebraska, which receives support from the UNL Office of Research and Innovation, and the Nebraska Research Initiative. This manuscript was partially written during the 2025 and 2026 editions of the Cedar Point Biological Station Writing Retreat. We also thank Dr. Pedro Pezzi for his comments in an earlier version of this manuscript.

## Supplementary tables

**Table S1.** Taxon sampling, including sample identifiers, dispersal phenotype classification. (FC, flight-capable; FL, flightless), number of Anchored Hybrid Enrichment (AHE) loci analyzed, mean depth and breadth of mitochondrial genome coverage recovered from off-target reads, and GenBank accession numbers for raw reads. Sequence data originally generated by Gray et al. (2020). Dispersal phenotype classifications compiled from Treidel et al. (2026).

## Supplementary figures

**Figure S1.** Maximum likelihood species tree inferred from the concatenated supermatrix of 560 Anchored Hybrid Enrichment loci using IQ-TREE under the GTR+G substitution model. Numbers at nodes indicate ultrafast bootstrap support values (1,000 replicates). Branch lengths are scaled in substitutions per site.

**Figure S2.** Gene Concordance Factors mapped onto the ASTRAL coalescent species tree. Values at each node indicate the percentage of informative gene trees that support that branch.

**Figure S3.** Site Concordance Factors mapped onto the ASTRAL coalescent species tree. Values at each node indicate the percentage of informative alignment sites supporting that branch, calculated across the full concatenated alignment.

**Figure S4.** Quartet Sampling results mapped onto the ASTRAL tree. Node labels show Quartet Concordance (QC), Quartet Differential (QD), and Quartet Informativeness (QI) scores, respectively. QC reflects the frequency with which sampled quartets support the focal branch; QD captures the symmetry of discordant quartets, with values near zero indicating asymmetric conflict consistent with introgression and values near 1 indicating symmetric conflict consistent with ILS; QI reflects the proportion of quartets that were phylogenetically informative at that node.

**Figure S5.** Internode Certainty All (ICA) scores mapped onto the ASTRAL coalescent species tree. ICA measures the degree to which a given bipartition is favored over all conflicting bipartitions in the set of gene trees. Values approaching 1 indicate strong and consistent gene tree support; values near zero or below indicate substantial conflict.

**Figure S6.** Maximum likelihood mitochondrial phylogeny inferred from mitochondrial loci assembled from off-target AHE reads, using IQ-TREE under the GTR+G substitution model. Numbers at nodes indicate ultrafast bootstrap support values (1,000 replicates). Branches are scaled in substitutions per site.

**Figure S7.** Phylogenetic networks inferred using SNaQ for models allowing zero through four reticulation events (h = 0–4). Each panel shows the best-scoring network topology for a given value of h. Curved branches indicate inferred hybridization events; numbers on hybrid edges indicate the inheritance probability (γ) at each hybrid node, representing the estimated proportion of the genome derived from each parental lineage.

**Figure S8.** Results from HyDe hybrid detection analyses. (A) Distribution of estimated admixture parameter values (γ) across the 4,912 triplet tests that detected significant levels of hybridization in North American *Gryllus*. (B) Permutation test evaluating the association between flight capability and the distribution of significant hybridization signals. The density curve shows the null distribution of the proportion of significant triplets containing at least one flight-capable species, generated from 10,000 random permutations of flight phenotype across taxa. The shaded region indicates the central 95% of the null distribution. The vertical line marks the observed proportion.

