## Supplementary figures and images for "A feedback between dispersal and hybridization may facilitate the repeated emergence of flight in field crickets"

### Figure S1

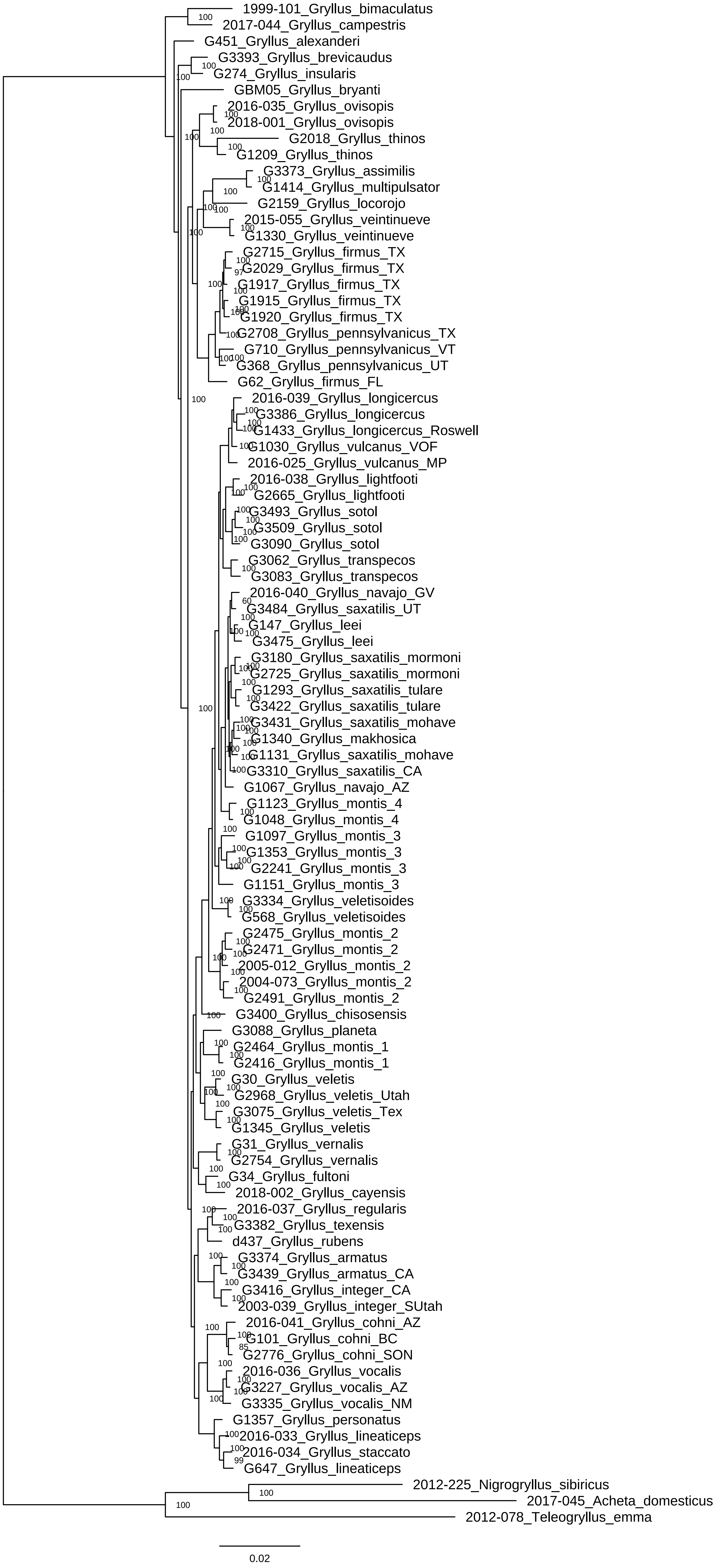

### Figure S2

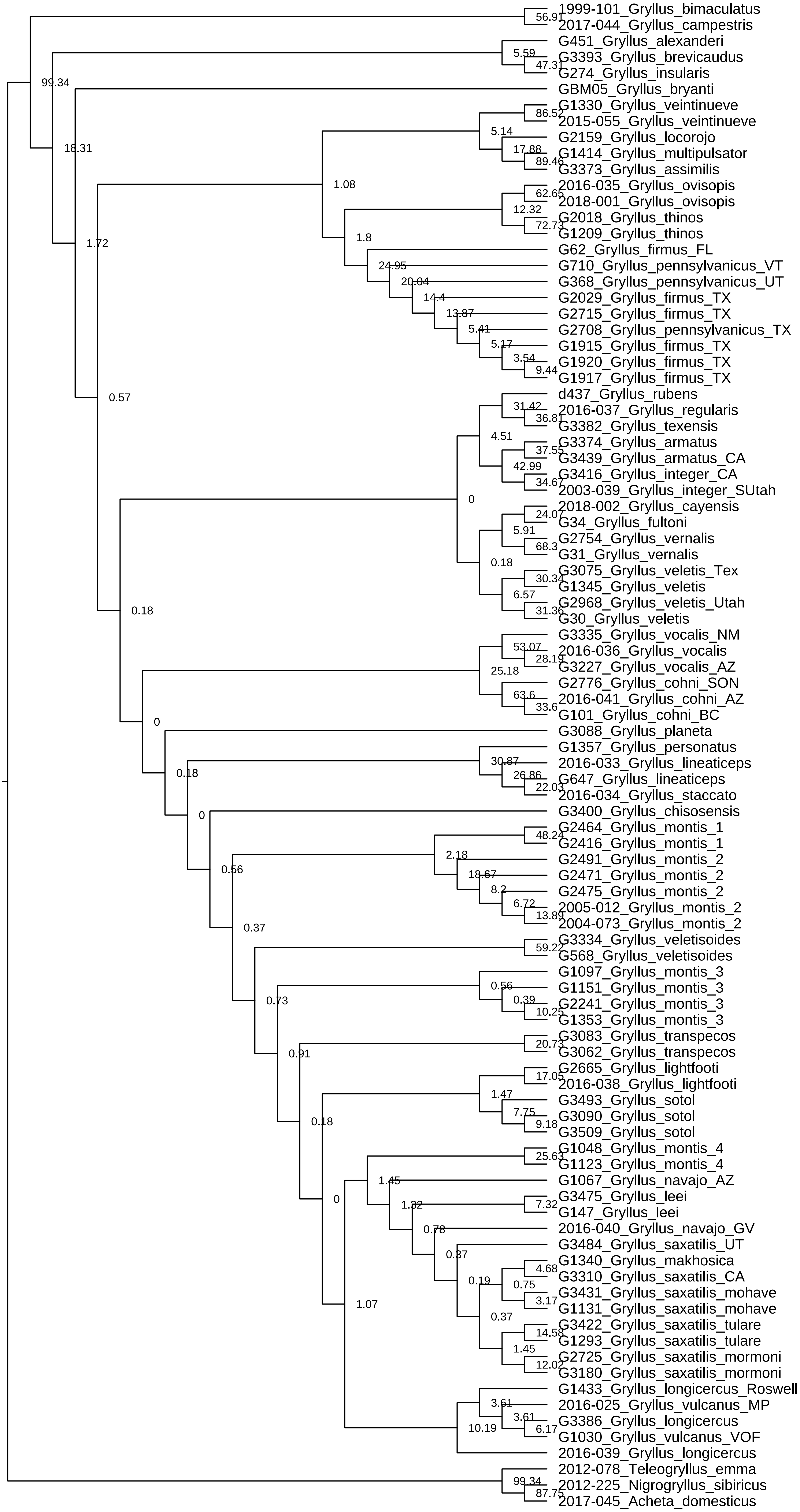

### Figure S3

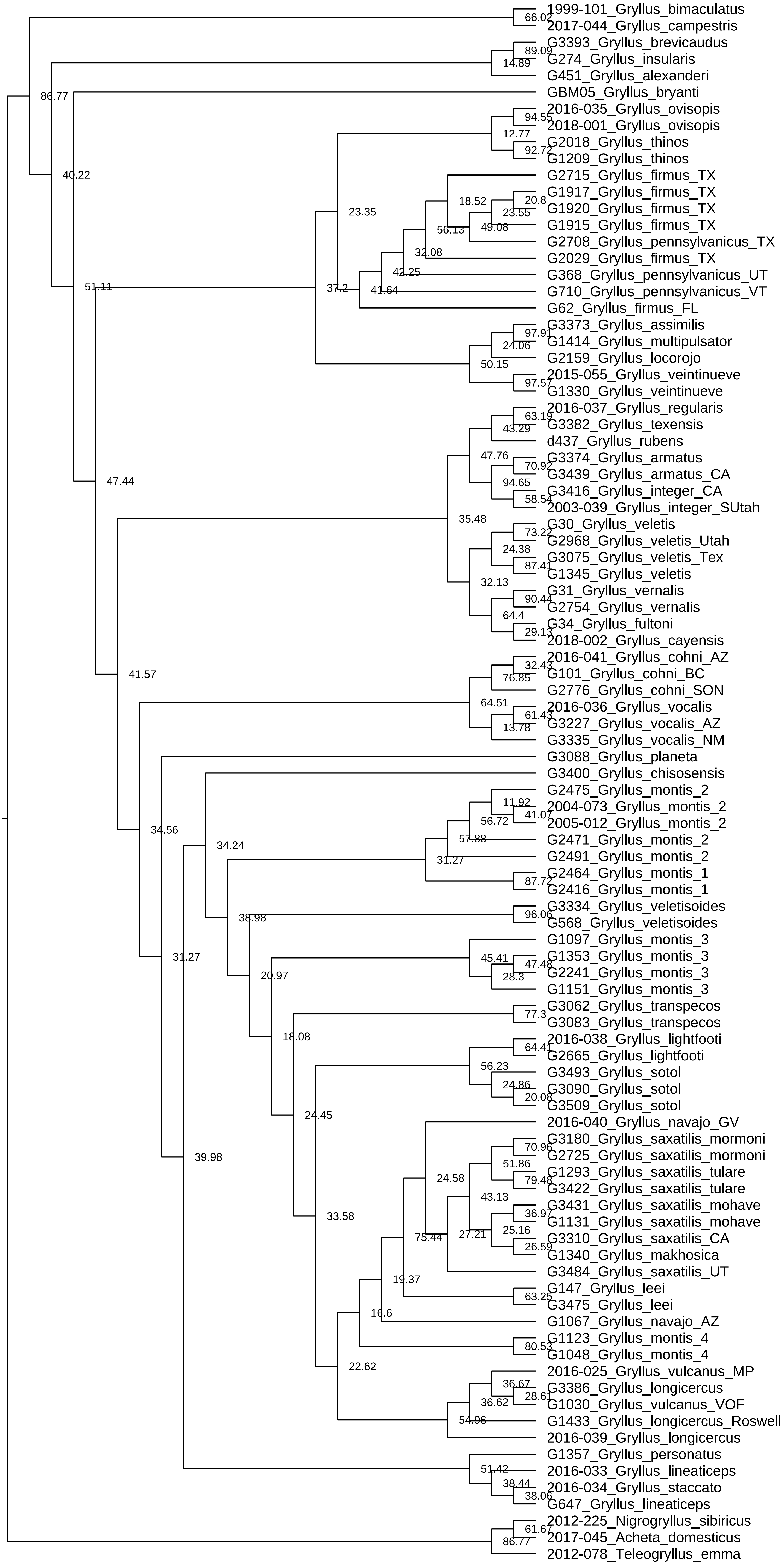

### Figure S4

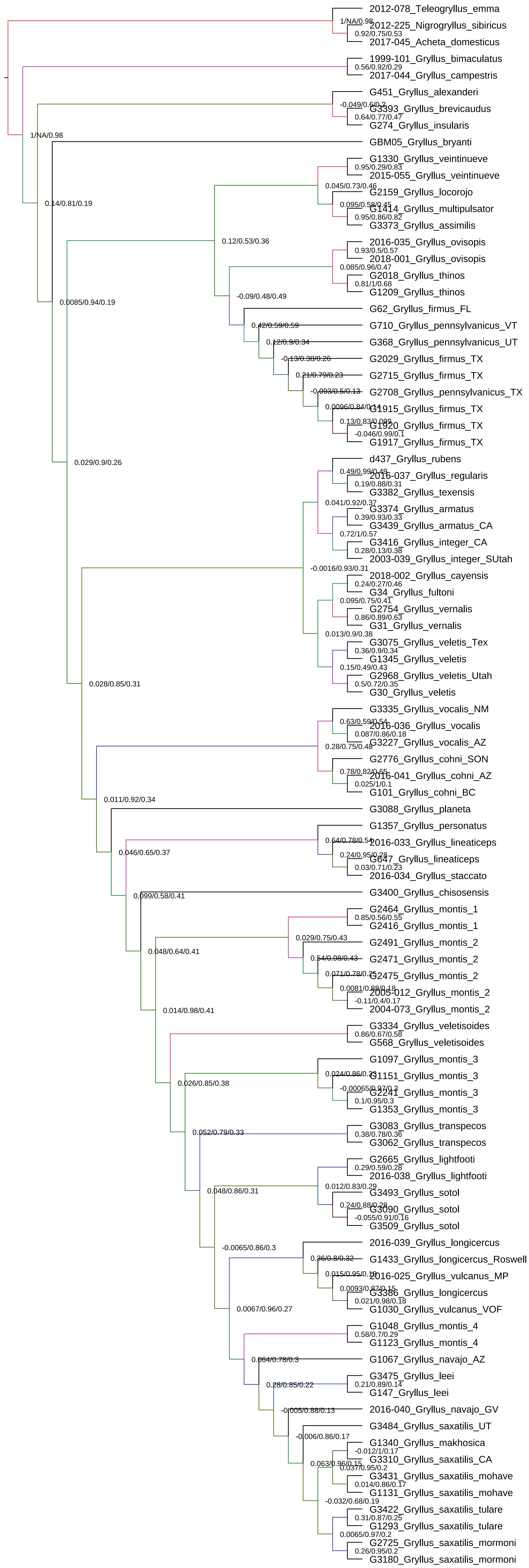

### Figure S5

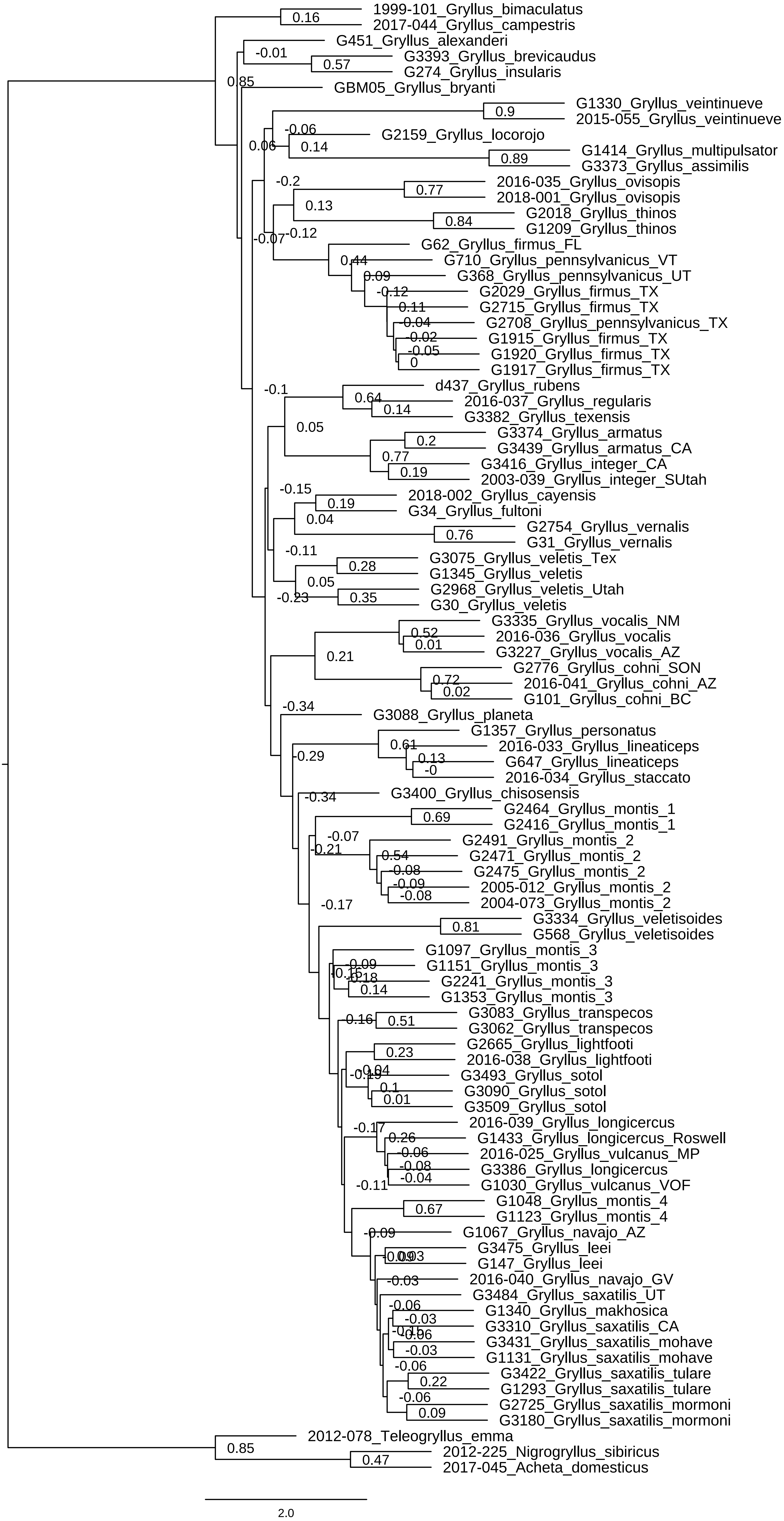

### Figure S6

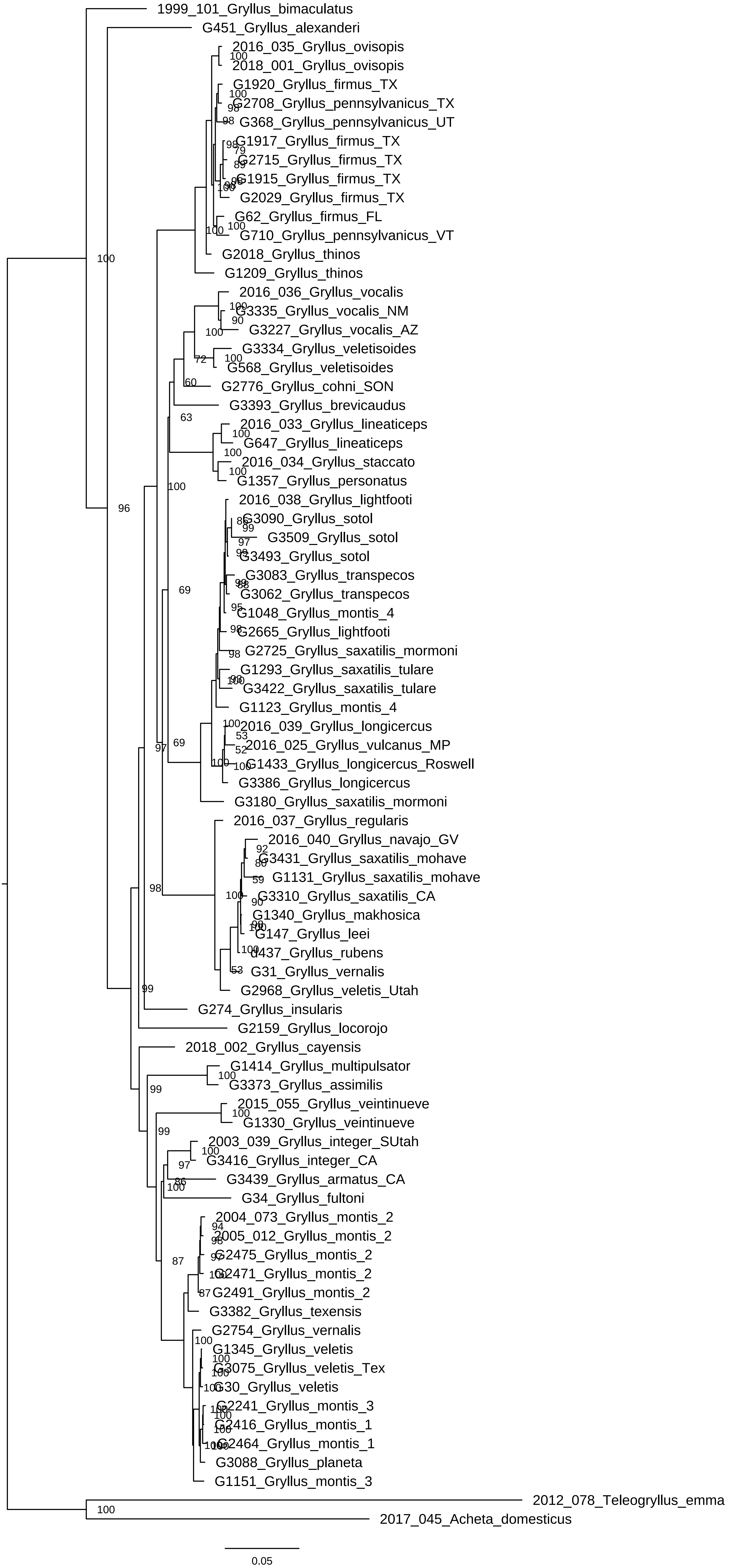

### Figure S7

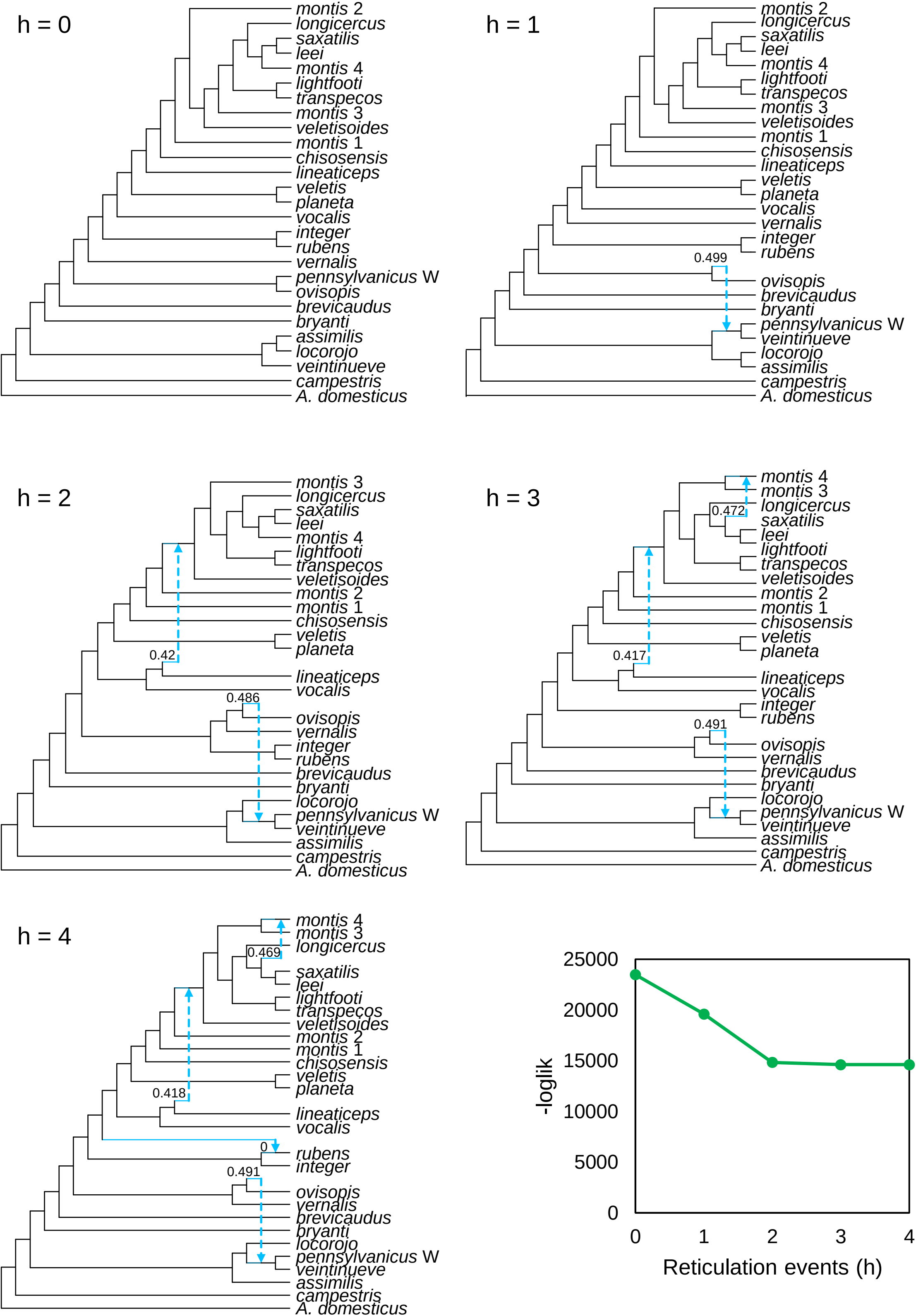

### Figure S8

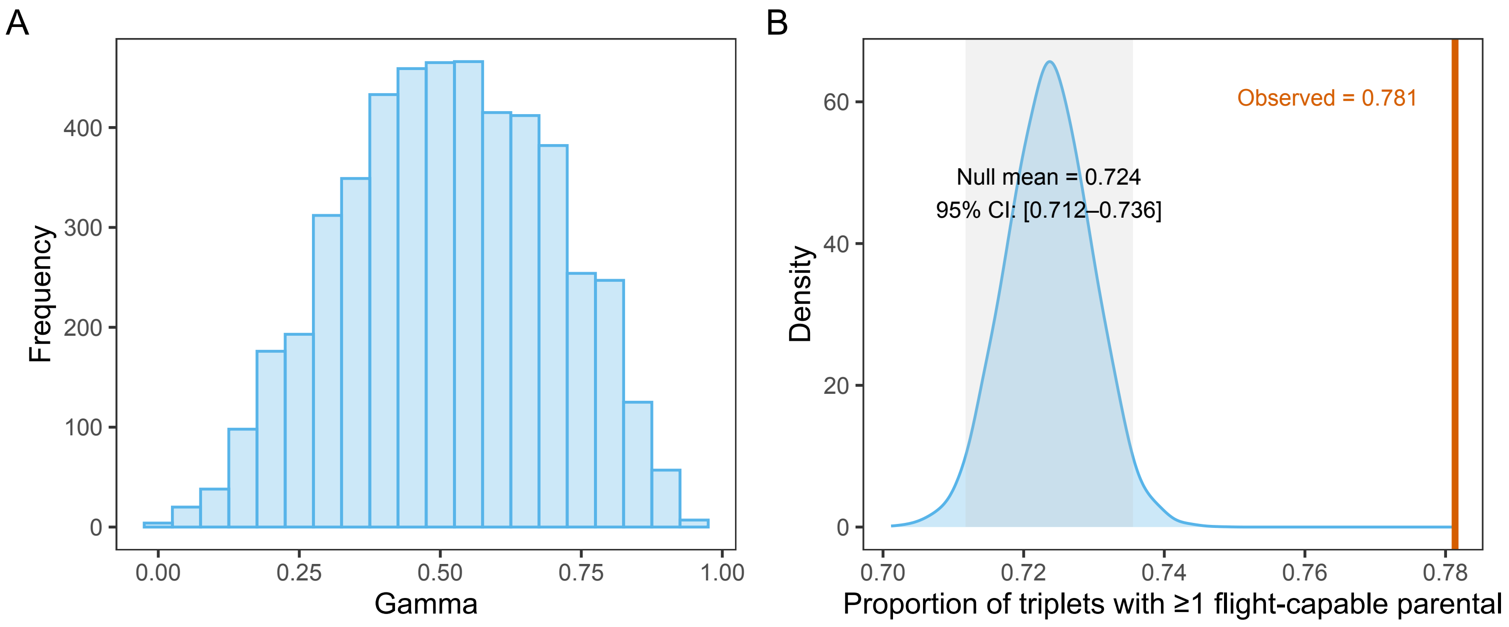
