## Supplementary material for "A feedback between dispersal and hybridization may facilitate the repeated emergence of flight in field crickets": Table S1

**Table S1.** Taxon sampling, including sample identifiers, dispersal phenotype classification (FC, flight-capable; FL, flightless), number of Anchored Hybrid Enrichment (AHE) loci analyzed, mean depth and breadth of mitochondrial genome coverage recovered from off-target reads, and GenBank accession numbers for raw reads. Sequence data originally generated by Gray et al. (2020). Dispersal phenotype classifications compiled from Treidel et al. (2026).

| **Species** | **ID** | **Flight** | **AHE loci** | **Depth** | **Breadth** | **Accession** |
|---|---|---|---|---|---|---|
| *Gryllus alexanderi* | G451 | FL | 528 | 34.3 | 92.4 | SRR40104099 |
| *Gryllus armatus* | G3374 | FC | 540 | 9.3 | 72.5 | SRR40104088 |
| *Gryllus armatus* | G3439 | FC | 539 | 26 | 95.1 | SRR40104077 |
| *Gryllus assimilis* | G3373 | FC | 539 | 9.7 | 77.2 | SRR40104066 |
| *Gryllus bimaculatus* | 1999-101 | FC | 395 | 35.6 | 99 | SRR40104055 |
| *Gryllus brevicaudus* | G3393 | FL | 534 | 31.4 | 93.9 | SRR40104044 |
| *Gryllus bryanti* | GBM05 * | FL | 528 | 6 | 71.8 | SRR40104033 |
| *Gryllus campestris* | 2017-044 * | FL | 538 | 8 | 68.4 | SRR40104022 |
| *Gryllus cayensis* | 2018-002 | FL | 536 | 15.7 | 92.1 | SRR40104011 |
| *Gryllus chisosensis* | G3400 * | FL | 536 | 9.1 | 74.2 | SRR40104098 |
| *Gryllus cohni* | 2016-041 * | FC | 539 | 7.8 | 70.5 | SRR40104097 |
| *Gryllus cohni* | G101 * | FC | 534 | 10.7 | 62.3 | SRR40104096 |
| *Gryllus cohni* | G2776 | FC | 530 | 25 | 97.5 | SRR40104095 |
| *Gryllus firmus* FL | G62 | FC | 542 | 132.3 | 97.8 | SRR40104094 |
| *Gryllus firmus* TX | G1915 | FC | 531 | 26.7 | 95.7 | SRR40104093 |
| *Gryllus firmus* TX | G1917 | FC | 531 | 35.8 | 97.1 | SRR40104092 |
| *Gryllus firmus* TX | G1920 | FC | 528 | 202.9 | 98.6 | SRR40104091 |
| *Gryllus firmus* TX | G2029 | FC | 528 | 43.4 | 98.9 | SRR40104090 |
| *Gryllus firmus* TX | G2715 | FC | 546 | 28.1 | 93.3 | SRR40104089 |
| *Gryllus fultoni* | G34 | FL | 522 | 157.8 | 98.8 | SRR40104087 |
| *Gryllus insularis* | G274 | FL | 534 | 16.2 | 82.8 | SRR40104086 |
| *Gryllus integer* | 2003-039 | FC | 540 | 71.8 | 96.4 | SRR40104085 |
| *Gryllus integer* | G3416 | FC | 539 | 41.8 | 93.8 | SRR40104084 |
| *Gryllus leei* | G147 | FL | 533 | 50 | 97.9 | SRR40104083 |
| *Gryllus leei* | G3475 * | FL | 503 | 6.6 | 63.2 | SRR40104082 |
| *Gryllus lightfooti* | 2016-038 | FC | 538 | 47.2 | 94.8 | SRR40104081 |
| *Gryllus lightfooti* | G2665 | FC | 534 | 15.4 | 89.3 | SRR40104080 |
| *Gryllus lineaticeps* | 2016-033 | FC | 543 | 71.1 | 93.8 | SRR40104079 |
| *Gryllus lineaticeps* | G647 | FC | 537 | 19.9 | 92 | SRR40104078 |
| *Gryllus locorojo* | G2159 | FC | 528 | 20.8 | 96.2 | SRR40104076 |
| *Gryllus longicercus* | 2016-039 | FC | 540 | 20.3 | 88 | SRR40104075 |
| *Gryllus longicercus* | G1433 | FC | 543 | 15.8 | 81.8 | SRR40104074 |
| *Gryllus longicercus* | G3386 | FC | 542 | 15.7 | 88.8 | SRR40104073 |
| *Gryllus makhosica* | G1340 | FL | 533 | 84.2 | 99.3 | SRR40104072 |
| *Gryllus montis* 1 | G2416 | FL | 533 | 58 | 99 | SRR40104062 |
| *Gryllus montis* 1 | G2464 | FL | 533 | 12 | 84.1 | SRR40104061 |
| *Gryllus montis* 2 | 2004-073 | FL | 533 | 416.5 | 98.6 | SRR40104071 |
| *Gryllus montis* 2 | 2005-012 | FL | 538 | 60.5 | 98.7 | SRR40104070 |
| *Gryllus montis* 2 | G2471 | FL | 525 | 23.9 | 97.2 | SRR40104060 |
| *Gryllus montis* 2 | G2475 | FL | 539 | 39.6 | 96.7 | SRR40104059 |

**Table S1** (continued from previous page)

| Species | ID | Flight | AHE loci | Depth | Breadth | Accession |
|---|---|---|---|---|---|---|
| *Gryllus montis* 2 | G2491 | FL | 528 | 25 | 98.1 | SRR40104058 |
| *Gryllus montis* 3 | G1097 * | FL | 535 | 12.5 | 68.6 | SRR40104068 |
| *Gryllus montis* 3 | G1151 | FL | 545 | 107.1 | 99.3 | SRR40104065 |
| *Gryllus montis* 3 | G1353 * | FL | 540 | 7 | 72.7 | SRR40104064 |
| *Gryllus montis* 3 | G2241 | FL | 530 | 39.4 | 99.4 | SRR40104063 |
| *Gryllus montis* 4 | G1048 | FL | 535 | 77.3 | 99 | SRR40104069 |
| *Gryllus montis* 4 | G1123 | FL | 534 | 33.3 | 93.5 | SRR40104067 |
| *Gryllus multipulsator* | G1414 | FC | 534 | 32.7 | 98.5 | SRR40104057 |
| *Gryllus navajo* | 2016-040 | FL | 538 | 14.1 | 86.1 | SRR40104056 |
| *Gryllus navajo* | G1067 * | FL | 387 | 3.8 | 38.7 | SRR40104054 |
| *Gryllus ovisopis* | 2016-035 | FL | 542 | 37.4 | 91 | SRR40104053 |
| *Gryllus ovisopis* | 2018-001 | FL | 531 | 51.3 | 98.4 | SRR40104052 |
| *Gryllus pennsylvanicus* | G710 | FC | 534 | 12.5 | 91.7 | SRR40104049 |
| *Gryllus pennsylvanicus* W | G2708 | FC | 533 | 29 | 98.5 | SRR40104051 |
| *Gryllus pennsylvanicus* W | G368 | FC | 539 | 9.7 | 84.9 | SRR40104050 |
| *Gryllus personatus* | G1357 | FC | 529 | 39.6 | 98.6 | SRR40104048 |
| *Gryllus planeta* | G3088 | FL | 542 | 19.5 | 90.5 | SRR40104047 |
| *Gryllus regularis* | 2016-037 | FC | 545 | 47.9 | 95.3 | SRR40104046 |
| *Gryllus rubens* | d437 | FC | 526 | 32.2 | 97.1 | SRR40104045 |
| *Gryllus saxatilis mohave* | G1131 | FL | 538 | 11.1 | 83.5 | SRR40104039 |
| *Gryllus saxatilis mohave* | G3431 | FL | 530 | 17.9 | 86 | SRR40104036 |
| *Gryllus saxatilis mormoni* | G2725 | FL | 531 | 16.8 | 89.5 | SRR40104043 |
| *Gryllus saxatilis mormoni* | G3180 | FL | 544 | 14.5 | 85 | SRR40104037 |
| *Gryllus saxatilis tulare* | G1293 | FL | 538 | 13 | 84 | SRR40104041 |
| *Gryllus saxatilis tulare* | G3422 | FL | 544 | 17.2 | 87.1 | SRR40104040 |
| *Gryllus saxatilis* | G3310 | FC | 527 | 18.5 | 97.8 | SRR40104042 |
| *Gryllus saxatilis* | G3484 * | FC | 540 | 9.2 | 71.8 | SRR40104038 |
| *Gryllus sotol* | G3090 | FL | 532 | 12.3 | 84.1 | SRR40104035 |
| *Gryllus sotol* | G3493 | FL | 527 | 34.5 | 98.6 | SRR40104034 |
| *Gryllus sotol* | G3509 | FL | 537 | 8.9 | 77.9 | SRR40104032 |
| *Gryllus staccato* | 2016-034 | FC | 535 | 62.9 | 96.8 | SRR40104031 |
| *Gryllus texensis* | G3382 | FC | 539 | 32.8 | 95.9 | SRR40104030 |
| *Gryllus thinos* | G1209 | FL | 529 | 56.9 | 98.7 | SRR40104029 |
| *Gryllus thinos* | G2018 | FL | 529 | 60.3 | 98.6 | SRR40104028 |
| *Gryllus transpecos* | G3062 | FL | 539 | 14.6 | 86.9 | SRR40104027 |
| *Gryllus transpecos* | G3083 | FL | 538 | 14.8 | 92.6 | SRR40104026 |
| *Gryllus veintinueve* | 2015-055 | FL | 528 | 28.1 | 98.2 | SRR40104025 |
| *Gryllus veintinueve* | G1330 | FL | 531 | 39.9 | 97.5 | SRR40104024 |
| *Gryllus veletis* | G1345 | FC | 544 | 41.5 | 96.4 | SRR40104023 |
| *Gryllus veletis* | G2968 | FC | 539 | 33 | 90.6 | SRR40104021 |
| *Gryllus veletis* | G30 | FC | 530 | 17.9 | 97.5 | SRR40104020 |
| *Gryllus veletis* | G3075 | FC | 542 | 28.9 | 89.9 | SRR40104019 |
| *Gryllus veletisoides* | G3334 | FL | 540 | 20.3 | 86.6 | SRR40104018 |
| *Gryllus veletisoides* | G568 | FL | 526 | 83 | 98.7 | SRR40104017 |
| *Gryllus vernalis* | G2754 | FL | 541 | 13.7 | 85.8 | SRR40104016 |

**Table S1** (continued from previous page)

| Species | ID | Flight | AHE loci | Depth | Breadth | Accession |
|---|---|---|---|---|---|---|
| *Gryllus vernalis* | G31 | FL | 526 | 23.8 | 98.2 | SRR40104015 |
| *Gryllus vocalis* | 2016-036 | FC | 541 | 28.6 | 94.6 | SRR40104014 |
| *Gryllus vocalis* | G3227 | FC | 537 | 11.7 | 87.3 | SRR40104013 |
| *Gryllus vocalis* | G3335 | FC | 543 | 14.8 | 86.7 | SRR40104012 |
| *Gryllus vulcanus* | 2016-025 | FL | 544 | 11.5 | 84.3 | SRR40104010 |
| *Gryllus vulcanus* | G1030 * | FL | 539 | 4.5 | 56.2 | SRR40104009 |
| *Acheta domesticus* | 2017-045 | NA | 480 | 10.9 | 73 | SRR40104100 |
| *Nigrogryllus sibiricus* | 2012-225 * | NA | 466 | 26.2 | 66.3 | SRR40104008 |
| *Teleogryllus emma* | 2012-078 | NA | 469 | 20.4 | 84.9 | SRR40104007 |

* Samples excluded from mitochondrial DNA analyses due to >50% missing data.
